# Assessing taxonomic metagenome profilers with OPAL

**DOI:** 10.1101/372680

**Authors:** Fernando Meyer, Andreas Bremges, Peter Belmann, Stefan Janssen, Alice C. McHardy, David Koslicki

## Abstract

Taxonomic metagenome profilers predict the presence and relative abundance of microorganisms from shotgun sequence samples of DNA isolated directly from a microbial community. Over the past years, there has been an explosive growth of software and algorithms for this task, resulting in a need for more systematic comparisons of these methods based on relevant performance criteria. Here, we present OPAL, a software package implementing commonly used performance metrics, including those of the first challenge of the Initiative for the Critical Assessment of Metagenome Interpretation (CAMI), together with convenient visualizations. In addition, OPAL implements diversity metrics from microbial ecology, as well as run time and memory efficiency measurements. By allowing users to customize the relative importance of metrics, OPAL facilitates in-depth performance comparisons, as well as the development of new methods and data analysis workflows. To demonstrate the application, we compared seven profilers on benchmark datasets of the first and second CAMI challenges using all metrics and performance measurements available in OPAL. The software is implemented in Python 3 and available under the Apache 2.0 license on GitHub (https://github.com/CAMI-challenge/OPAL).

**Author summary:** There are many computational approaches for inferring the presence and relative abundance of taxa (i.e. taxonomic profiling) from shotgun metagenome samples of microbial communities, making systematic performance evaluations a very important task. However, there has yet to be introduced a computational framework in which profiler performances can be compared. This delays method development and applied studies, as researchers need to implement their own custom evaluation frameworks. Here, we present OPAL, a software package that facilitates standardized comparisons of taxonomic metagenome profilers. It implements a variety of performance metrics frequently employed in microbiome research, including runtime and memory usage, and generates comparison reports and visualizations. OPAL thus facilitates and accelerates benchmarking of taxonomic profiling techniques on ground truth data. This enables researchers to arrive at informed decisions about which computational techniques to use for specific datasets and research questions.

## Introduction

Taxonomic metagenome profilers predict the taxonomic identities and relative abundances of microorganisms of a microbial community from shotgun sequence samples. In contrast to taxonomic binning, profiling does not result in assignments for individual sequences, but derives a summary of the presence and relative abundance of different taxa in microbial community. In some use-cases, such as pathogen identification for clinical diagnostics, accurate determination of the presence or absence of a particular taxon is important, while for comparative studies, such as quantifying the dynamics of a microbial community over an ecological gradient, accurately determining relative abundances of taxa is paramount.

Given the variety of use-cases, it is important to understand the benefits and drawbacks of the particular taxonomic profiler for different applications. While there has been much effort in developing taxonomic profiling methods [1–12], only recently have community efforts arisen to perform unbiased comparisons of such techniques and assess their strengths and weaknesses [13, 14]. Critical obstacles to such comparisons have been a lack of consensus on performance metrics and output formats by the community, as different taxonomic profilers report their results in a variety of formats and interested parties had to implement their own metrics for comparisons.

Here, we describe OPAL (Open-community Profiling Assessment tooL), a framework that directly addresses these issues. OPAL aggregates the results of multiple taxonomic profilers for one or more benchmark datasets, computes relevant metrics for different applications on them, and then presents the relative strengths and weaknesses of different tools in intuitive graphics. OPAL leverages the emerging standardized output format recently developed by the CAMI consortium [13, 15] to represent a taxonomic profile and which has been implemented for a variety of popular taxonomic profilers [2, 4–10, 12]. OPAL can also use the popular BIOM format [16]. The metrics that OPAL computes range from simple presence-absence metrics to more sophisticated comparative metrics such as UniFrac [17] and diversity metrics. The resulting metrics are displayed in graphics viewable in a browser and allow a user to dynamically rank taxonomic profilers based on the combination of metrics of their choice.

Similar efforts to provide comparative frameworks have recently been made for genome binners of metagenome samples (AMBER [18]) and metagenomic assemblers (QUAST [19, 20]). OPAL augments these efforts by addressing the issue of comparing and assessing taxonomic profilers. OPAL will assist future systematic benchmarking efforts. It will aid method developers to rapidly assess how their implemented taxonomic profilers perform in comparison to other techniques, and facilitate assessing profiler performance characteristics, such as clarifying when and where tool performance degrades (eg. performance at particular taxonomic ranks). Importantly, OPAL will help to decide which profiler is best suited to analyze particular datasets and biological research questions, which vary widely, depending on nature of the sampled microbial community, experimental setup and sequencing technology used [21].

## Methods

### Inputs

OPAL accepts as inputs one or several taxonomic profiles and benchmarks them at different taxonomic ranks against a given taxonomic gold standard profile.

Both the predicted and gold standard taxonomic profiles may contain information for multiple samples, such as for a time series, technical or biological replicates. A gold standard taxonomic profile can, for instance, be created with the CAMISIM metagenome simulator [21, 22]. The taxonomic profiles can be either in the Bioboxes profiling format [15, 23] or the BIOM (Biological Observation Matrix) format [16]. Examples are provided in the OPAL GitHub repository [24].

### Metrics and accompanying visualizations

OPAL calculates a range of relevant metrics commonly used in the field [13] for one or more taxonomic profiles of a given dataset by comparing to a gold standard taxonomic profile. Below, we give formal definitions of all metrics, together with an explanation of their biological meaning.

#### Preliminaries

For *r*, a particular taxonomic rank (or simply rank), let *x_r_* be the true bacterial relative abundances at rank *r* given by the gold standard. That is, *x_r_* is a vector indexed by all taxa at rank *r*, where entry (*x_r_*)_*i*_ is the relative abundance of taxon *i* in the sampled microbial community at rank *r*. With 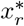 we denote the vector of predicted bacterial relative abundances at rank *r*. Accordingly, 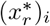 is the predicted relative abundance of taxon *i* at rank *r*.

By default, OPAL normalizes all (predicted) abundances prior to computing metrics, such that the sum of all abundances equals 1 at each rank, i.e. Ʃ_*i*_(*x_r_*)_*i*_ = 1 and 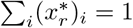. This is to avoid any bias towards profiling software that makes fewer predictions, say, for only 50% of the sample.

#### Assessing the presence or absence of taxa

The purity and completeness of taxonomic predictions are common measures for assessing profiling quality [25]. They assess how well a profiler correctly identifies the presence and absence of taxa in a sampled microbial community without considering how well their relative abundances were inferred. This can be relevant, for example, in an emergency situation in clinical diagnostics, when searching for a pathogen in a metagenomic sample taken from patient material. To define these measures, let the support of the vector *x_r_* be

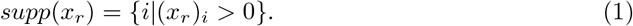

That is, *supp*(*x_r_*) is the set of indices of the taxa at rank *r* present in the sample. Analogously,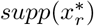 is the set of indices of the taxa at rank *r* predicted to be in the sample. For each rank *r*, we define the true positives *TP_r_*, false positives *FP_r_*, and false negatives *FN_r_*, respectively, as

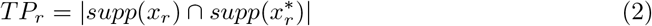

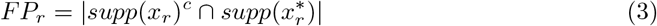

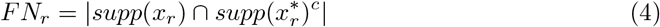

where *supp*(*x_r_*)^*c*^ and 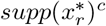 are the complement of the respective support vectors, and, thus, give the indices of the taxa at rank *r* absent or predicted as absent in the sample. Specifically, *TP_r_* and *FP_r_* are the number of taxa correctly and incorrectly predicted as present in the sample, respectively, and *FN_r_* is the number of taxa incorrectly predicted as being absent in the sample.

The **purity *p_r_*** at rank *r*, also known as precision or specificity, is the ratio of taxa correctly predicted as present in the sample and all predicted taxa at that rank. For each rank *r*, the purity is computed as

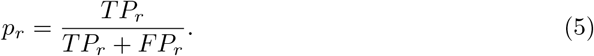

The **completeness *s_r_*** at rank *r*, also known as recall or sensitivity, is the ratio of taxa correctly predicted as present and all taxa present in the sample at that rank. For each taxonomic rank *r*, the completeness is computed as

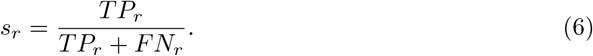

Purity and completeness range from 0 (worst) to 1 (best).

We combine purity and completeness into a single metric by computing their harmonic average, also known as the **F1 score**. It is defined for each rank *r* as

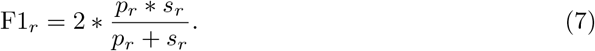

The F1 score ranges from 0 to 1, being closer to 0 if at least one of the metrics purity or completeness has a low value, and closer to 1 if both the purity and completeness are high.

The **Jaccard index *J*** is a common metric to determine the percentage of organisms common to two populations or samples. We define it as an indicator of similarity between the sets of true and predicted taxa at each rank by computing the ratio of the number of taxa in the intersection of these sets to the number of taxa in their union. Formally, it is computed for each rank as

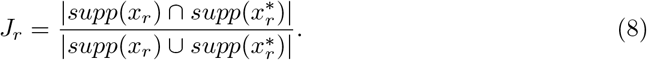

The Jaccard index ranges from 0 (complete dissimilarity) to 1 (complete overlap).

#### Abundance estimates

The next category of metrics for assessing profiling quality not only considers whether taxa was predicted as present or absent in the sample, but also their abundances.

The L1 norm measures the accuracy of reconstructing the relative abundance of taxa in a sample at rank *r*. The L1 norm is given by

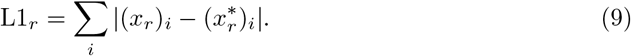

The **L1 norm** thus gives the total error between the true and predicted abundances of the taxa at rank *r*. It ranges from 0 to 2, where 0 indicates perfect reconstruction of the relative abundances of organisms in a sample and 2 indicates totally incorrect reconstruction of relative abundances.

Another metric, the **Bray-Curtis distance *d_r_***, is derived from the L1 norm by dividing the sum of the absolute pairwise differences of taxa abundances by the sums of all abundances at the given rank. This bounds the Bray-Curtis distance between 0 and 1. For each rank *r*, it defined as

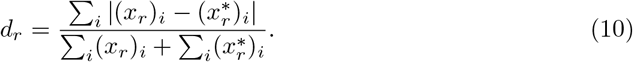

The **weighted UniFrac distance** is a tree-based measure of taxonomic similarity of microbial communities [17] measuring the similarity between true and predicted abundances. Instead of a phylogenetic tree as in [17], we use a taxonomic tree with nodes restricted to eight major ranks and store the true and predicted abundances on the appropriate nodes. In summary, the UniFrac distance is the total amount of predicted abundances that must be moved (along the edges of the taxonomic tree, with all branch lengths here set to 1) to cause them to overlap with the true relative abundances. We use the EMDUnifrac implementation of the UniFrac distance [26–28]. A low UniFrac distance indicates that a taxonomic profiling algorithm gives a prediction that is taxonomically similar to the actual profile of the sample. The weighted UniFrac distance ranges between 0 and twice the height of the taxonomic tree used. Because each level of the tree represents one of the ranks superkingdom, phylum, class, order, family, genus, species, and strain, the maximum weighted UniFrac distance is 16.

The **unweighted UniFrac distance** is similar to the weighted UniFrac distance, but instead of storing the *relative abundances* for the appropriate nodes, a 1 is placed on the node if the profile indicates a non-zero relative abundance at that node and a 0 otherwise. Hence, it can be considered a measure of how well (in terms of taxonomic similarity) a profiler correctly identified the presence and absence of taxa in a sample. The maximum unweighted UniFrac distance is equal to

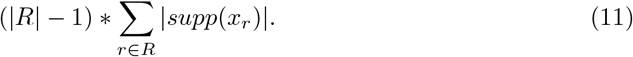

where *R* is the set of all taxonomic ranks.

#### Alpha diversity metrics

Unlike the metrics above, alpha diversity metrics are computed from a single profile of (predicted) abundances at each rank, without a comparison to e.g. a gold standard profile. Alpha diversity metrics summarize the variety (or richness) and distribution of taxa present in a profile [29] and, among other uses, is commonly used to observe global shifts in community structure as a result of some environmental parameter [30–33].

The simplest alpha diversity metric is the number of taxa present in a given environment. We measure this at each rank individually for a given profiler, allowing a comparison to the underlying gold standard. For a given profile *x_r_* (or 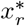), we denote the number of taxa at rank *r* as *S_r_* = |*supp*(*x_r_*)|.

As a measure of diversity also considering the relative taxon abundances, we combine *S_r_* and all abundances (*x_r_*)_*i*_ (or 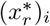) using the **Shannon diversity index *H_r_*** [34]. For each rank *r*, it is calculated as

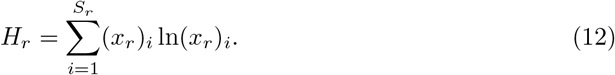

*H_r_* ranges from 0 to ln(*S_r_*), where ln(*S_r_*) represents the maximal possible diversity, with all taxa being evenly represented. We note that the Shannon diversity index traditionally assumes that all taxa are represented in the sample. However, because some profilers may not predict abundances for all taxa, we ignore such taxa in the sum (where 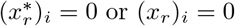)

While *H_r_* accounts for diversity and evenness, the **Shannon equitability index *E_r_*** is a measure of evenness. It is a normalized form of the Shannon diversity index obtained by dividing *H_r_* by its maximum value ln(*S_r_*), i.e.

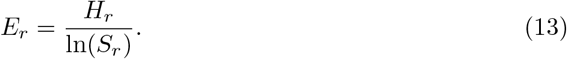

Thus, *E_r_* ranges from 0 to 1 with 1 indicating complete evenness.

#### Beta diversity metrics

In contrast to alpha diversity, beta diversity metrics give an indication of taxa distribution similarity between a pair of profiles [29]. If beta diversity is small, not only is the diversity similar between the profiles, but the actual *distribution* of relative abundances between profiles are similar. To compare the similarity of beta diversity predictions for each profiler versus the gold standard, we display the following information in a scatter plot. Each point corresponds to a pair of input samples with the x-coordinate being the Bray-Curtis distance between the taxonomic profilers predictions on the pair of samples. The y-coordinate is the Bray-Curtis distance between the gold standards corresponding to the pair of samples. The closer this scatter plot is to the line *y* = *x*, the more closely the taxonomic profiler results in taxa distributions similar to the gold standard. These plots are shown at each taxonomic rank.

#### Rankings

To indicate a global sense of relative performance, we also rank profilers by their relative performance across each sample, taxonomic rank, and metric. In particular, each profiler is assigned a score for its performance for each metric within a taxonomic rank and sample. The best performing profiler gets score 0, the second best, 1, and so on. These scores are then added over the taxonomic ranks and samples to produce a single score per metric for each profiler. Also, an overall score of each profiler is computed by summing up all its scores per metric. The resulting scores are displayed in an interactive table of an HTML page, with a row per profiler, a column per metric, and an additional column for the overall scores. The columns can be sorted by the user and, therefore, yield a ranking of the profilers over all metrics or for a specific one. Optionally, the overall score of each profiler can be computed by summing up its score per metric in a weighted fashion, i.e., a user can interactively select custom weighting on the HTML page, depending on the combination of metrics that most suits their needs. The default weight of each metric is 1 and can vary between 0 and 10, in steps of 0.1. For example, if a user is interested in profilers that are highly precise and accurately reconstruct the exact relative abundance of predicted taxa, they can emphasize purity and L1 norm (e.g. giving each weight 3) over UniFrac error and completeness (e.g. giving each weight 1). The resulting rankings are dynamically updated in real time and graphically presented to the user.

#### Output and visualizations

OPAL outputs the assessment of the predictions of multiple profilers in several formats: flat files, tables (per profiling program, taxonomic rank, and in tidy format [35]), plots, and in an interactive HTML visualization. An example page is available at [36]. The visualizations created include:

- **Absolute performance plots:** To visually compare the relative performance of multiple profilers, spider plots (also known as radar plots) of completeness and purity are created, with the spokes labeled with the corresponding profiler name. At least three profilers are required for these plots. The completeness and purity metrics are shown as colored lines connecting the spokes, with the scale on the spokes indicating the value of the error metric. One such spider plot is created at each taxonomic rank to give an indication of performance versus rank.
- **Relative performance plots:** Similarly, spider plots are created for the completeness, purity, false positives, weighted UniFrac, and L1 norm for three or more profilers. Since the values of these metrics have very different scales, they are each normalized by the maximum value attained by any input profiler. Hence, these plots indicate the relative performance of each profiler with respect to the different metrics. For example, one profiler having the largest value of the purity metric would indicate that, amongst the compared profilers, it is the most precise (without indicating what the exact value of the purity metric is). These plots are also shown at each taxonomic rank.
- **Shannon equitability:** The Shannon equitability index is plotted against taxonomic ranks for each input profile along with the gold standard. This results in a visual indication of how closely a taxonomic profile reflects the actual alpha diversity of the gold standard.
- **Bray-Curtis distances:** For each profiler, a scatter plot of Bray-Curtis distances is created to compare the similarity of beta diversity of the profiler predictions versus the gold standard. For details, see the section above on beta diversity metrics.
- **Ranking:** In a bar chart shown on the created HTML page, each bar corresponds to the sum of scores obtained by a profiler as a result of its ranking for the metrics completeness, purity, L1 norm, and weighted UniFrac over all major taxonomic ranks. The bar chart is dynamically updated in real time according to the weight assigned to each metric by the user. For details of the computation of the scores, see the above section on rankings.
- **Taxa proportions:** For each taxonomic rank, a stacked bar chart shows the taxa proportions in each sample of the gold standard, with each bar corresponding to a sample and each color to a taxon. This gives a visual indication of the taxa abundances and variations among the samples. On the HTML page, the user may opt to see a legend of the colors and corresponding taxa. The legend is only optionally displayed since the number of taxa can vary between a few superkingdoms to hundreds or thousands of species or strains, and these cannot all be reasonably displayed on a single image.
- **Rarefaction and accumulation curves:** A plot simultaneously shows rarefaction and accumulation curves for all the major taxonomic ranks. To ease the visualization at different ranks, another plot shows the curves in logarithmic scale with base 10.

## Results and discussion

To demonstrate an application, we evaluated taxonomic profilers on two datasets. First, we evaluated taxonomic profiling submissions to the first CAMI challenge [13] on the dataset with the highest microbial complexity in the challenge. We will call this dataset CAMI I HC for short. This is a simulated time series benchmark dataset with 5 samples, each with size 15 Gbp, and a total of 596 genomes. It includes Bacteria, Archaea, and high-copy circular elements (plasmids and viruses) with substantial real and simulated strain-level diversity. We reproduce and extend the results for this dataset from [13] with alpha and beta diversity metrics implemented in OPAL and measure the run time and memory usage of profiling methods. The second dataset that we evaluated taxonomic profilers on were the short-read data of a new *practice* dataset of the second CAMI challenge (CAMI II MG, for short). This consists of 64 samples with a total size of 320 Gbp, and was simulated from taxonomic profiles for microbial communities from the guts of different mice [21]. This resulted in the inclusion of 791 genomes as *meta-community* members from public databases. The samples in both CAMI I HC and CAMI II MG are paired-end 150 bp Illumina reads and are available at [37, 38].

To visualize the taxonomic composition and properties of these datasets, we produced plots of the taxa proportions at all major taxonomic ranks for all samples with OPAL (Figs A and B in S1 Supporting Information for CAMI II MG and CAMI I HC, respectively) and calculated rarefaction curves (Fig C in S1 Supporting Information).

The assessed profilers were CommonKmers (an early version of MetaPalette) [2], a combination of Quikr [8], ARK [9], and SEK [10] (abbreviated Quikr), TIPP 2.0.0 [12], Metaphlan 2.2.0 [5], MetaPhyler 1.25 [6], mOTU 1.1 [7], and FOCUS (CAMI version) [4]. The software parameters used are available in Table A in S1 Supporting Information. The reference databases used by each profiler precede the release of the genomes used for generating the first CAMI challenge datasets. Thus the metagenomic information of the CAMI I HC dataset was completely new for these profilers and at different taxonomic distances to available reference genomes, differently from the metagenome data of the CAMI II MG practice dataset. The profilers were run as Bioboxes docker containers on a computer with an Intel Xeon E5-4650 v4 CPU (virtualized to 16 CPU cores, 1 thread per core) and 512 GB of main memory. Metaphlan was the fastest method on CAMI II MG with a run time of 12.5 hours, whereas on CAMI I HC, Metaphlan and Quikr were the fastest methods, requiring roughly the same execution time of 2.12 hours (Fig 1, Table B in S1 Supporting Information). mOTU was the most memory efficient method on both datasets (1.19 GB of maximum main memory usage), closely followed by Metaphlan (1.44 and 1.66 GB maximum main memory usage on CAMI II MG and CAMI I HC respectively).

**Fig 1.**
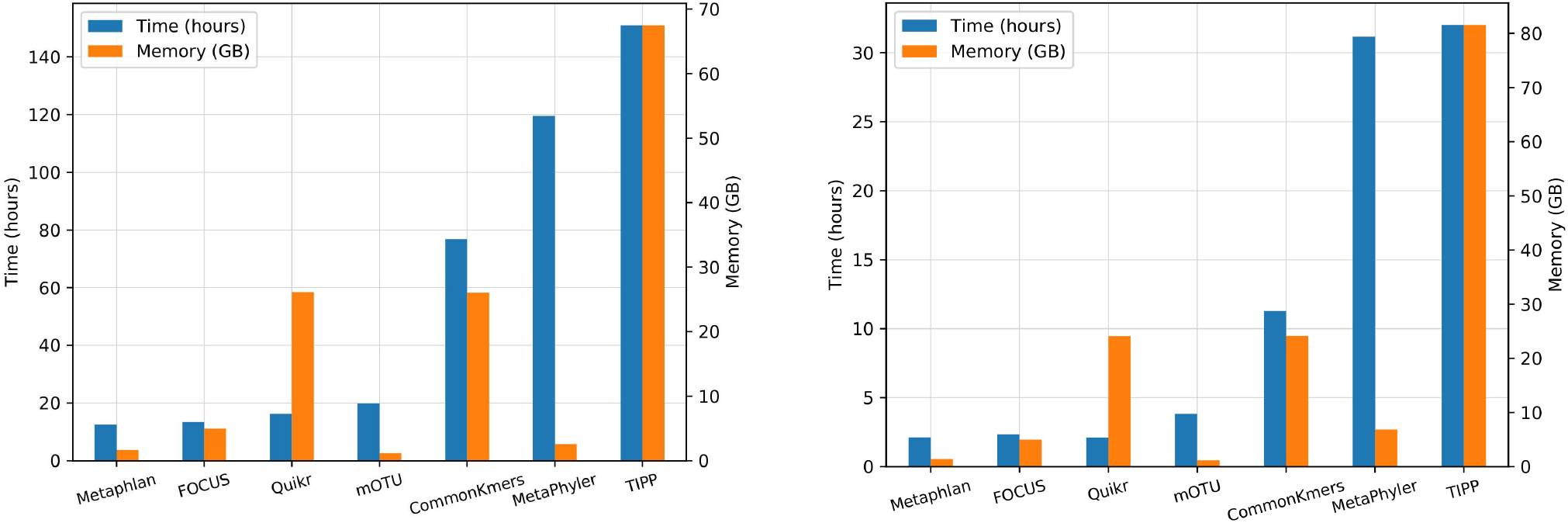
Computing efficiency. Run time in hours and maximum main memory usage in gigabytes required the by profilers to process the CAMI II mouse gut (left) and the CAMI I high complexity (right) datasets.

On the CAMI I HC data, Quikr, TIPP, and MetaPhyler, in this order, achieved the overall highest completeness (Figs D(a-c) and E-G(h-n) in S1 Supporting Information). However, these profilers obtained the lowest purity. In this metric, CommonKmers and Metaphlan performed best. In terms of the F1 score, computed from completeness and purity, Metaphlan was the best method. This indicates that Metaphlan performed particularly well in determining the presence or absence of taxa. However, it could not accurately predict their relative abundances, as indicated by the high L1 norm error. In this metric, MetaPhyler did well, followed by FOCUS and CommonKmers.

When ranking methods over all taxonomic ranks using completeness, purity, L1 norm, and weighted UniFrac with equal weights (Figs D(c) and H(b) in S1 Supporting Information), TIPP performed best with total score 184. TIPP ranked second for completeness and weighted UniFrac (scores 31 and 5, respectively), third for L1 norm (score 52), and only for purity it did not do so well and was ranked fifth (score 96). When considering the performance of the profilers at different taxonomic ranks, we found that most profilers performed well until the family level. For example, TIPP and MetaPhyler achieved a 0.92 completeness at the family level, but this decreased to 0.43 at the genus level. Similarly, the purity of CommonKmers decreased from 0.96 at the family level to 0.77 and 0.08 at the genus and species levels, respectively.

In terms of alpha diversity, no profiler estimated taxon counts well. Most programs overestimated diversity at all taxonomic ranks. Quikr, FOCUS, and CommonKmers predicted taxon abundances that better reflect the the Shannon equitability of the gold standard. However, Quikr, mOTU, and TIPP made no predictions at the strain level. The predicted abundance distributions of CommonKmers and mOTU across all samples at the species level best reflect the gold standard, as visualized with the scatter plots of Bray-Curtis distances (Fig J in S1 Supporting Information). Taken together, the OPAL results fully reproduce the results from [13], where performance was summarized in three categories of profilers: profilers that correctly predicted relative abundances, profilers with high purity, and those with high completeness. OPAL extends the overall performance view by providing analysis of computing efficiency and microbial diversity predictors.

On the CAMI II MG data, Metaphlan obtained the overall best ranking over all taxonomic ranks, using the equally weighted metrics completeness, purity, L1 norm, and weighted UniFrac (Fig 2(d) and Fig H(a) in S1 Supporting Information). MetaPhyler achieved the highest completeness at most taxonomic ranks, followed by TIPP and Metaphlan (Figs E-G(a-g) in S1 Supporting Information), whereas CommonKmers achieved the highest completeness at the species level (Fig 2(c)). Metaphlan was not only among the profilers with the highest completeness, but it also maintained a high purity throughout all taxonomic ranks, with only a small decrease from genus (0.94) to species (0.89). This can be explained by a high coverage of CAMI II MG by the reference genomes used by Metaphlan. It also contrasts with the results in [13], showing that a profiler can be precise while achieving a relative high completeness, but with this being very dependent on the input data. Metaphlan also predicted taxon distributions across the samples well. MetaPhyler and TIPP could not identify well differences in taxa abundances for the samples and tended to predict similar abundances, which is reflected in many points in the plots being above the line *x* = *y* (Fig 3(b-h)).

**Figure 2.**
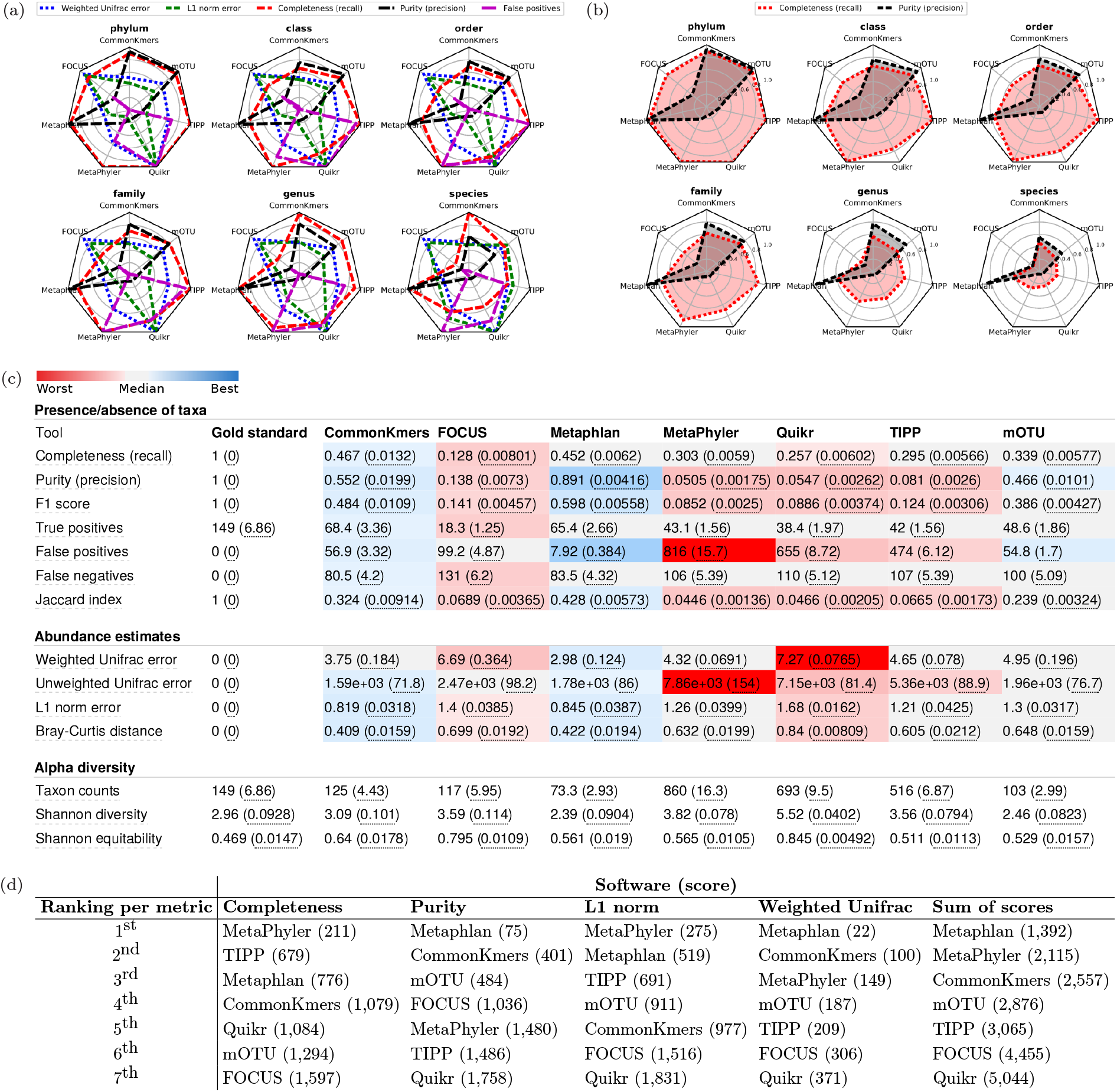
Assessment results on the CAMI II mouse gut dataset. (a) Relative performance plots with results for the metrics: weighted UniFrac, L1 norm, completeness, purity, and number of false positives at different taxonomic ranks. The values of the metrics in these plots are normalized by the maximum value attained by any profiler at a certain rank. (b) Absolute performance plots with results for the metrics completeness and recall, ranging between 0 and 1. (c) Results at the species level for all computed metrics, as output by OPAL in the produced HTML page. The values are averaged over the results for all 64 samples of the dataset, with the standard error being shown in parentheses. The colors indicate the quality of the prediction by a profiler with respect to a metric, from best (dark blue) to worst (dark red). (d) Rankings of the profilers according to their performance and scores for different metrics computed over all samples and taxonomic ranks.

**Figure 3.**
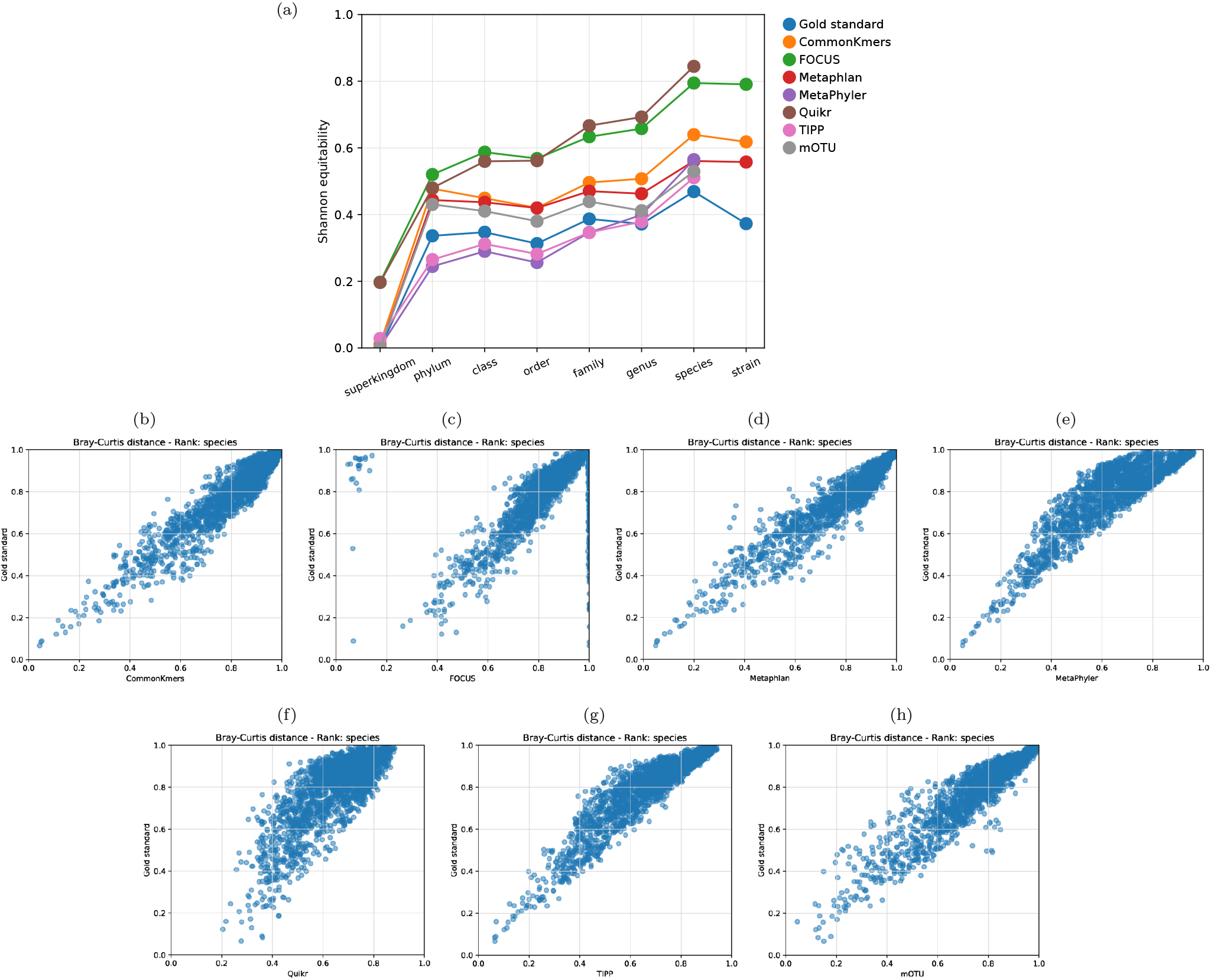
Examples of alpha and beta diversity plots from the results on the CAMI II mouse gut dataset. (a) Shannon equitability at different taxonomic ranks as a measure of alpha diversity. The closer the Shannon equitability of the predicted profile by a method to the gold standard, the better it reflects the actual alpha diversity in the gold standard in terms of evenness of the taxa abundances. (b-h) Scatter plots of Bray-Curtis distances visualizing beta diversity at the species level. For each profiling method and plot, a point corresponds to the Bray-Curtis distance between the abundance predictions for a pair of input samples by the method (x-axis) and the Bray-Curtis distance computed for the gold standard for the same pair of samples (y-axis). The closer a point is to the line *x* = *y*, the more similar the predicted taxa distributions are to the gold standard.

In terms of alpha diversity, Metaphlan, CommonKmers, and mOTU predicted taxon counts similar to the gold standard for most taxonomic ranks, whereas the other profilers mostly overestimated the counts. On the other hand, TIPP, MetaPhyler, and mOTU predicted taxon abundances that more closely reflect their evenness, i.e., Shannon equitability, in the gold standard (Fig 3(a) and Fig I(a-b) in S1 Supporting Information). As on the CAMI I HC data, Quikr, mOTU, and TIPP made no strain-level predictions on this dataset.

## Conclusions

OPAL facilitates performance assessment and interpretation for taxonomic profilers using shotgun metagenome datasets as input. It implements commonly used performance metrics, including diversity metrics from microbial ecology, and outputs the assessment results in a convenient HTML page, in tables, and plots. By providing rankings and the possibility to give different weights to the metrics, OPAL enables the selection of the best profiler suitable for a researchers particular biological interest. In addition, computational efficiency results that OPAL returns can guide users on the choice of a profiler under time and memory constraints. We plan to continually extend the metrics and visualizations of OPAL according to community requirements and suggestions.

We used OPAL to analyze the CAMI I HC data, demonstrating how it enables reproduction of the results of this study [13]. We also used it for the analysis of a new large dataset, the CAMI II MG data, revealing consistency across many metrics and softwares analysed, and also a few striking differences. Specifically, while on the CAMI I HC data Quikr had the highest completeness by a wide margin, on the CAMI II MG data MetaPhyler performed best with this metric and Quikr was among the least complete profiling tools. Similarly, the Metaphlan results changed from the lowest to the highest weighted UniFrac score. Results such as these indicate the importance of choosing a program suitable for the particular properties of the microbial community analyzed and considering variables such as the availability of reference genome sequences of closely related organisms to those in the sample. Given the wide variety of environments from which metagenome data are obtained, this further demonstrates the relevance of OPAL.

## Supporting information

**S1 Supporting Information. Additional tables and figures.**

